## Supplemental figures and tables for "DIP/Dpr interactions and the evolutionary design of specificity in protein families"

| Protein | Dpr4 | Dpr6 | Dpr10 | Dpr11 | DIP-γ | DIP-α |
|---|---|---|---|---|---|---|
| Uniprot ID | Q59DX6 | M9PC40 | M9PF17 | Q8MRE6 | Q9VAR6 | Q9W4R3 |
| 1 | D74 | N102 | N82 | N146 | K67 | G70 |
| 2 | R75 | K103 | K83 | K147 | N68 | Y71 |
| 3 | A76 | T104 | T84 | S148 | K69 | R72 |
| 4 | S78 | S106 | A86 | S150 | G71 | G74 |
| 5 | I80 | I108 | I88 | I152 | L73 | L76 |
| 6 | R81 | R109 | R89 | R153 | R74 | K77 |
| 7 | K82 | H110 | H90 | L154 | A75 | A78 |
| 8 | R83 | R111 | R91 | R155 | S76 | D79 |
| 9 | D84 | D112 | D92 | D156 | D77 | T80 |
| 10 | L85 | I113 | L93 | G157 | Q78 | K81 |
| 11 | H86 | H114 | H94 | H158 | T79 | A82 |
| 12 | I87 | I115 | I95 | I159 | V80 | I83 |
| 13 | V90 | V118 | V98 | V162 | L83 | I85 |
| 14 | G91 | G119 | G99 | D163 | Q84 | H86 |
| 15 | L93 | Y121 | Y101 | A165 | R86 | N89 |
| 16 | Y95 | Y123 | Y103 | F167 | V88 | I91 |
| 17 | T96 | T124 | T104 | I168 | T89 | T92 |
| 18 | N97 | S125 | T105 | A169 | H90 | H93 |
| 19 | D98 | D126 | D106 | D170 | N91 | N94 |
| 20 | Q99 | Q127 | Q107 | Q171 | A92 | P95 |
| 21 | R100 | R128 | R108 | R172 | R93 | R96 |
| 22 | Q120 | Q148 | Q128 | Q191 | R112 | S115 |
| 23 | R122 | R150 | R130 | R193 | S114 | E117 |
| 24 | D123 | D151 | D131 | D194 | D116 | D118 |
| 25 | E128 | E156 | E136 | E199 | M120 | M123 |
| 26 | Q130 | Q158 | Q138 | Q201 | Q122 | Q125 |
| 27 | S132 | S160 | S140 | S203 | N124 | N127 |
| 28 | T133 | T161 | T141 | T204 | T125 | T128 |
| 29 | E134 | Q162 | Q142 | E205 | S126 | D129 |
| 30 | P135 | P163 | P143 | P206 | P127 | P130 |
| 31 | K136 | V164 | V144 | K207 | M128 | M131 |
| 32 | S138 | S166 | S146 | S209 | K130 | S133 |
| 33 | G140 | F168 | S148 | R211 | V132 | I135 |

Family-wide numbering of interfacial positions

UniProt numbering

**Supplementary Table 1:** The correspondence between protein specific and family-wide numbering. Family wide-numbering (as defined in Fig. 2A) is compared to numbering of amino acid residues of Dpr4, Dpr6, Dpr10, Dpr11, DIP-γ, and DIP-α in UniProt database.

Binding affinity change

-3.00 ΔΔG, kcal/mol +3.00

stabilizing destabilizing

| Interaction | Mutation (a) | Mutation (b) | See footnote | Experiment | FoldX | mCSM | BeAtMusic | Mutabind | Rosetta flex | BindProfX |
|---|---|---|---|---|---|---|---|---|---|---|
| Dpr6/DIP-α | Dpr6 H110K | Dpr6 H7K | (c) | 1.91 | 0.46 | 0.91 | 0.61 | 1.45 | 0.83 | 1.59 |
| | DIP-α K81Q | DIP-α K10Q | (d) | 1.31 | 1.06 | 0.88 | 0.89 | 1.39 | -0.03 | 0.58 |
| | DIP-α S133D | DIP-α S32D | (e) | 0.21 | 0.52 | 0.59 | 0.51 | 0.43 | 0.17 | 0.00 |
| | DIP-α G74S | DIP-α G4S | (e) | 1.01 | 0.86 | 1.41 | 0.64 | 0.59 | 1.21 | 2.48 |
| | DIP-α A82T | DIP-α A11T | (d) | 0.85 | 1.90 | 0.54 | 0.55 | 0.90 | 1.06 | 1.08 |
| | DIP-α N94D | DIP-α N19D | (d) | -0.45 | 0.12 | 0.52 | 0.33 | 1.03 | 0.06 | 0.89 |
| | DIP-α G74A | DIP-α G4A | (d) | -0.46 | -0.28 | 1.12 | 0.10 | 0.20 | 0.80 | 2.11 |
| | DIP-α G74L | DIP-α G4L | (d) | 1.35 | 6.72 | 1.47 | -0.10 | 2.29 | 4.63 | 2.51 |
| | DIP-α A78K | DIP-α A7K | (d) | -0.95 | -0.26 | 1.01 | 1.30 | 0.58 | -0.19 | 1.12 |
| | DIP-α I91A | DIP-α I16A | (d) | 2.18 | 1.79 | 0.72 | 1.80 | 1.77 | 0.97 | 0.76 |
| Dpr4/DIP-η | Dpr4 K82H | Dpr4 K7H | (c) | 0.12 | 0.12 | 0.08 | 0.19 | 0.70 | -0.34 | 0.87 |
| DIP-α/DIP-α | DIP-α K81Q | DIP-α K10Q | (d) | -0.12 | 1.71 | 3.06 | 0.92 | 1.87 | -0.06 | 0.58 |
| | DIP-α G74S | DIP-α G4S | (e) | 0.51 | 0.07 | 3.04 | 0.04 | 1.03 | 1.76 | 0.00 |
| | DIP-α S133D | DIP-α S32D | (e) | 0.70 | 0.06 | 0.84 | 0.60 | 0.94 | 0.46 | 0.00 |
| Dpr10/DIP-α | Dpr10 Q138D | Dpr10 Q26D | (d) | 1.36 | 3.82 | -0.47 | 1.13 | 2.51 | 2.20 | 0.83 |
| | DIP-α K81Q | DIP-α K10Q | (d) | 1.85 | 1.49 | 1.01 | 0.75 | 1.05 | 0.78 | 0.58 |
| | DIP-α D129S | DIP-α D29S | (e) | 0.16 | -0.85 | 1.46 | 1.13 | 0.61 | -0.24 | 1.61 |
| | DIP-α G74S | DIP-α G4S | (e) | 0.65 | -1.07 | 1.26 | 0.38 | 0.41 | 1.01 | 2.48 |
| | DIP-α S133D | DIP-α S32D | (e) | 0.32 | 0.26 | 0.50 | 0.47 | 0.36 | 0.20 | 0.00 |
| | DIP-α I91A | DIP-α I16A | (d) | 2.11 | 1.27 | 0.84 | 1.97 | 1.44 | 0.87 | 0.76 |
| | DIP-α G74L | DIP-α G4L | (d) | 1.88 | 6.55 | 1.27 | -0.29 | 1.92 | 3.36 | 2.51 |
| | DIP-α A82T | DIP-α A11T | (d) | 1.33 | 0.87 | 0.30 | 0.36 | 0.76 | 0.33 | 1.08 |
| | DIP-α N94D | DIP-α N19D | (d) | -0.13 | 0.11 | 0.21 | 0.24 | 0.79 | 0.34 | 0.89 |
| | DIP-α G74A | DIP-α G4A | (d) | -0.42 | -0.25 | 0.96 | -0.12 | 0.17 | 0.80 | 2.11 |
| | DIP-α A78K | DIP-α A7K | (d) | -1.09 | -0.47 | 1.19 | 1.17 | 0.48 | -0.09 | 1.12 |
| | | | | PCC | 0.54 | -0.13 | 0.19 | 0.61 | 0.49 | 0.05 |
| | | | | RMSE | 1.38 | 1.21 | 1.56 | 0.92 | 1.41 | 1.40 |

(a) Protein specific residue numbering of all the mutants as in Uniprot database (see Supplementary Table 1 for details)
(b) Family-wide residue numbering used for interfacial positions throughout the paper (as indicated in Fig. 2)
(c) Value calculated based on previously published SPR binding affinity ($K_D$) measurements for wild-type (WT) and single mutant (MT) proteins using the following formula: $\Delta\Delta G = RT \ln (K_{D(MT)} / K_{D(WT)})$, Cosmanescu et al. Neuron (2018).
(d) Value calculated based on SPR binding affinity ($K_D$) measurements for wild-type (WT) and single mutant (MT) proteins using the following formula: $\Delta\Delta G = RT \ln (K_{D(MT)} / K_{D(WT)})$. The supporting SPR data can be found in Supplementary Fig. 1.
(e) Value for the indicated mutation (X) was calculated in the context of a background mutation (DIP-α K81Q). Mutation X is >12Å away from the background mutation (K81Q), allowing us to compare effect of the computed single X mutation to the experimental value, which is calculated based on SPR binding affinity ($K_D$) measurements for mutant DIP-α K81Q (context) and double mutant DIP-α K81Q X (X) proteins using the following formula: $\Delta\Delta G = RT \ln (K_{D(X)} / K_{D(context)})$.

Pearson correlation coefficient (PCC) and lower root mean square error (RMSE) are calculated for every method (FoldX, mCSM, BeAtMusic, Mutabind, Rosetta flex, BindProfX) based on comparison of the theoretically calculated 25 datapoints with the set of 25 experimental values (Experiment).

**Supplementary Table 2: Performance of computational tools to predict changes in binding affinity for a set of DIP and Dpr mutants.** Higher Pearson correlation coefficient (PCC) and lower root mean square error (RMSE) reflect better agreement with experimental values (see methods). ΔΔG values are color-coded as indicated in the scale accompanying the data. Calculations were performed on the following PDB structures: 5EO9, 6EG0, 6EFY, 6NRQ (chains C,D).

A

| Protein | Mutation | $K_D$ (SPR) to a cognate **DIP-α** partner, μM | ΔΔG (SPR), kcal $mol^{-1}$ | ΔΔG (FoldX), kcal $mol^{-1}$ |
|---|---|---|---|---|
| **Dpr6** | wild-type | 2.3 | 0 | 0 |
| **Dpr6** | H7K {H110K} | 57.5 | 1.91 | 0.46 ± 0.20 |
| **Dpr6** | H7K V31K {H110K V164K} | >300 | >2.89 | 1.24 ± 0.64 |

B

| Protein | Mutation | $K_D$ (SPR) to non-cognate **DIP-α** partner, μM |
|---|---|---|
| **Dpr4** | wild-type | >1000 |
| **Dpr4** | K7H {K82H} | >400 |
| **Dpr4** | K7H K31V {K82H K136V} | 44.9 |

C

| Protein | Mutation | $K_D$ (SPR) to cognate **DIP-η** partner, μM | ΔΔG (SPR), kcal $mol^{-1}$ | ΔΔG (FoldX), kcal $mol^{-1}$ |
|---|---|---|---|---|
| **Dpr4** | wild-type | 83.7 | 0 | 0 |
| **Dpr4** | K7H {K82H} | 50.9 | -0.29 | -0.05 ± 0.03 |
| **Dpr4** | K7H K31V {K82H K136V} | >400 | 0.93 | 2.02 ± 0.09 |

D

| Protein | Mutation | $K_D$ (SPR) to non-cognate **DIP-η** partner, μM |
|---|---|---|
| **Dpr6** | wild-type | >1000 |
| **Dpr6** | H7K {H110K} | >1000 |
| **Dpr6** | H7K V31K {H110K V164K} | >700 |

**Supplementary Table 3: Comparison of experimental SPR measurements with FoldX data validating predictions of negative constraints in blue and green DIP/Dpr subfamilies.** Effect of mutations on **(A)** DIP-α/Dpr6 binding in SPR; **(B)** DIP-α/Dpr4 binding in SPR; **(C)** DIP-η/Dpr4 binding in SPR; **(D)** DIP-η/Dpr6 binding in SPR. DIPs and Dprs are color-coded according to the subgroups they are in Figure 1B. Family-wide residue numbering is followed by UniProt numbering (in curly brackets) for each mutant. FoldX calculations were performed on cognate complexes.

| | Dpr11,15,16,17 | Dpr6,10 | Dpr8,9,21 | Dpr13,14,18,19,20 | Dpr12 | Dpr1,2,3,4,5,7 | Dpr7 |
|---|---|---|---|---|---|---|---|
| DIP-γ | | Dpr: 7, 14, 16, 29, 31<br>DIP: 9, 10, 16 | Dpr: 7, 12, 14, 16, 29, 31<br>DIP: 9, 11, 15 | Dpr: 14, 16, 29, 31<br>DIP: 7, 9, 10, 11, 18, 29 | Dpr: 13, 14, 16, 29, 31<br>DIP: 7, 9, 11 | Dpr: 14, 16<br>DIP: 7, 18 | Dpr: 13, 14, 16, 27<br>DIP: 5, 7, 16, 29, 33 |
| DIP-α | Dpr: 10, 29, 31<br>DIP: 6, 9, 11, 15, 22 | | Dpr: 12, 29, 31<br>DIP: 4, 5 | Dpr: 15, 31<br>DIP: 4, 5, 7, 9, 10, 18, 29 | Dpr: 7, 10, 13, 18, 20, 31<br>DIP: 4, 7, 9, 10, 11, 29 | Dpr: 7, 29, 31<br>DIP: 4, 18, 23 | Dpr: 2, 7, 13, 27, 29, 31<br>DIP: 3, 4, 6, 7 |
| DIP-β,λ | Dpr: 12, 29, 31<br>DIP: 6, 9, 11, 15 | Dpr: 12, 29, 31<br>DIP: 5 | | Dpr: 29<br>DIP: 5, 7, 9, 10, 29 | Dpr: 10, 12, 13, 31<br>DIP: 7, 9, 10, 11, 29 | Dpr: 12, 29, 31<br>DIP: 5, 7 | Dpr: 12, 13, 27, 29, 31<br>DIP: 5, 6, 7 |
| DIP-ε,ζ | Dpr: -<br>DIP: 6, 9, 10, 11, 15, 31 | Dpr: 31<br>DIP: 10, 13, 30, 31 | Dpr: 29<br>DIP: 5, 10, 13, 31 | | Dpr: 13, 31<br>DIP: 5, 7, 9, 11, 31, 32, 33 | Dpr: 33<br>DIP: 10, 13, 18, 31 | Dpr: 13, 33<br>DIP: 5, 6, 10, 13, 18, 31, 32, 33 |
| DIP-δ | Dpr: 10, 14, 16, 17, 29, 31<br>DIP: 6, 15, 19, 22 | Dpr: 7, 10, 18, 20, 31<br>DIP: 5, 10 | Dpr: 7, 10, 12, 20, 31<br>DIP: 19 | Dpr: 29, 31<br>DIP: 5, 7, 10, 29 | | Dpr: 7, 10, 29, 31<br>DIP: 5, 9 | Dpr: 7, 10, 29, 31, 32<br>DIP: 5, 6, 18, 33 |
| DIP-ι,θ,η | Dpr: 10<br>DIP: 9, 15 | Dpr: 31<br>DIP: 10 | Dpr: 1, 12, 29, 31<br>DIP: 5, 11, 20 | Dpr: 31<br>DIP: 5, 7, 9, 10, 11, 29 | Dpr: 10, 13, 31<br>DIP: 9, 11, 23, 29 | | Dpr: -<br>DIP: 5, 18 |
| DIP-κ | Dpr: 2, 10, 27, 32<br>DIP: 5, 9, 15, 22 | Dpr: 2, 27, 29, 31, 32<br>DIP: 10 | Dpr: 12, 27, 29, 31, 32<br>DIP: 5, 11, 15 | Dpr: 29, 31<br>DIP: 5, 7, 9, 10, 11, 29 | Dpr: 10, 27, 31, 32<br>DIP: 5, 9, 11, 23, 29 | Dpr: 32<br>DIP: 5, 22 | |

**Supplementary Table 4: Negative constraints predicted between non-cognate DIP/Dpr subfamilies.** Interfacial positions destabilizing non-cognate binding are listed on both Dpr and DIP side in off-diagonal elements. Interfacial positions correspond to family-wide numbering presented in Fig.2 and compared to Uniprot numbering in Supplementary Table 1.

A

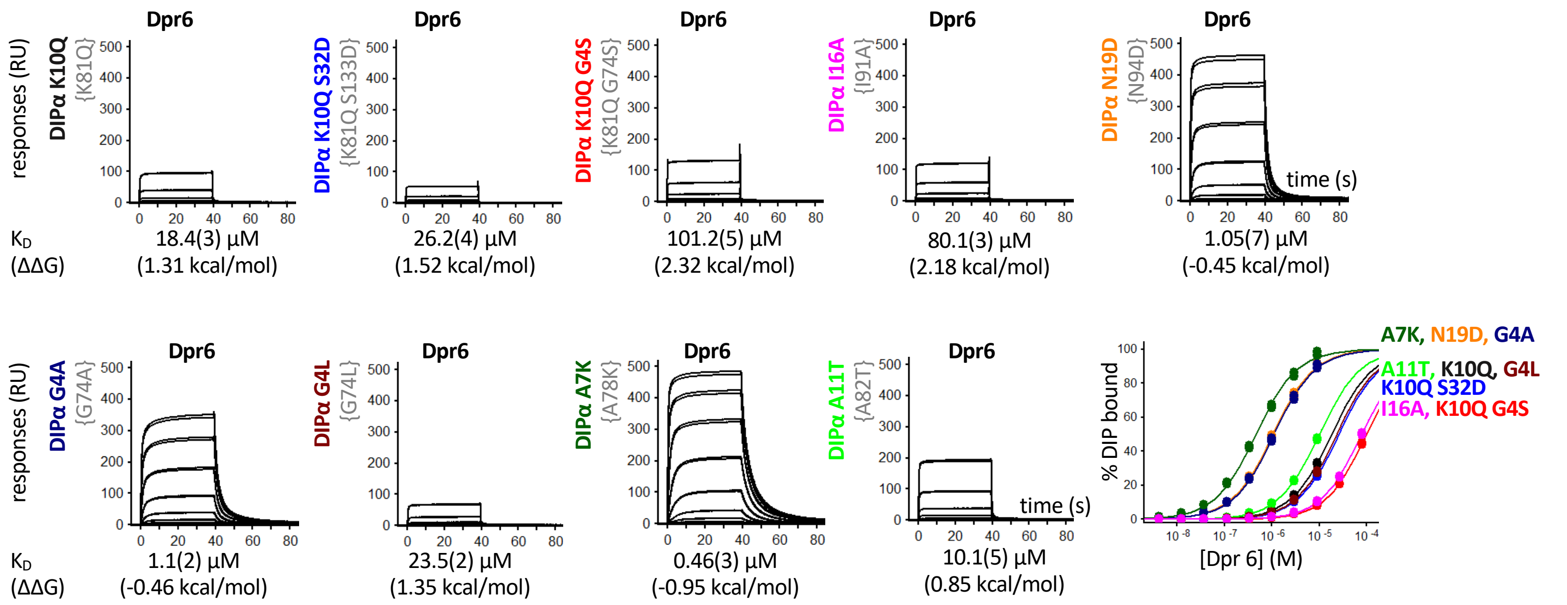


B

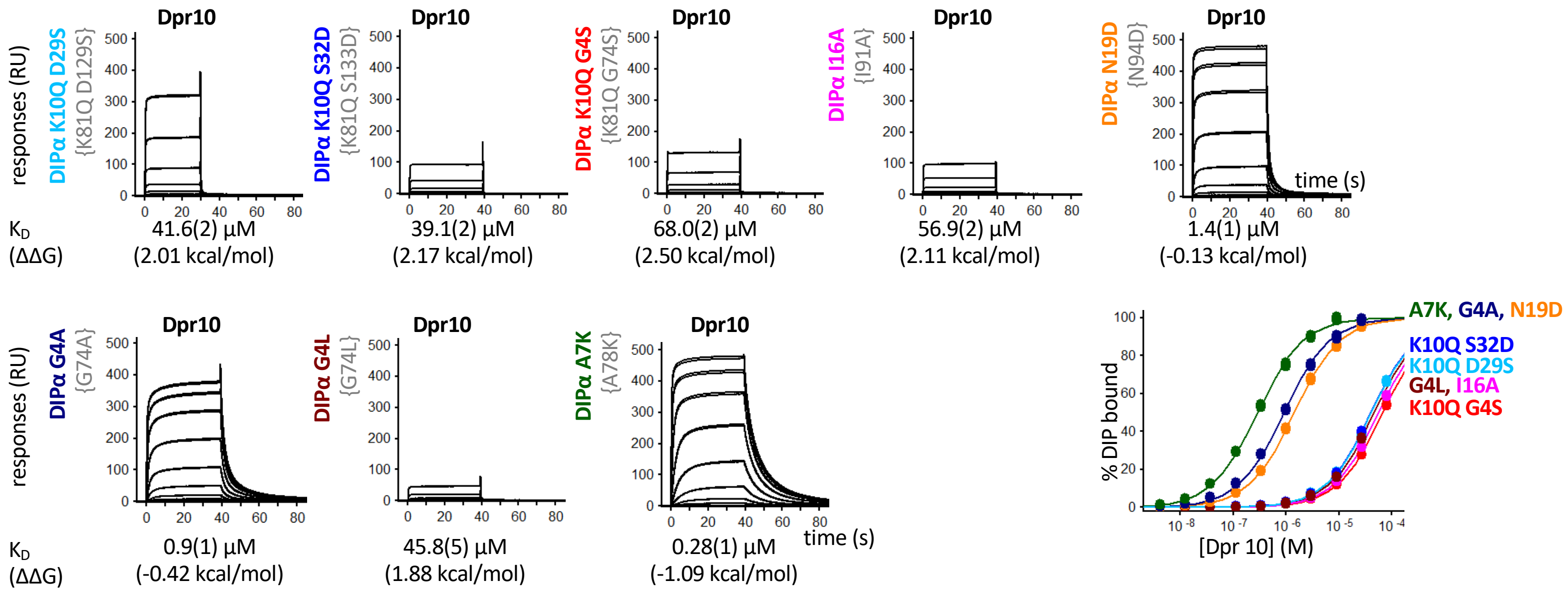


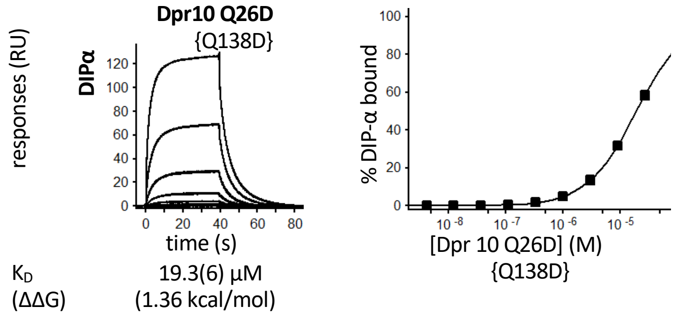


**Supplementary Fig. 1**: SPR and AUC experiments of mutants presented in Supplementary Table 2. Effect of mutations on **(A)** DIP-α/Dpr6 binding in SPR; **(B)** DIP-α/Dpr10 binding in SPR; and **(C)** DIP-α/DIP-α binding in AUC. Analyte Dpr proteins were flown over the chip surface with immobilized DIPs (A and B). $K_D$ values of DIP/Dpr interactions are given below the SPR sensorgrams. The number in parenthesis represents the fitting error in the last significant figure, in µM, for a single experiment with an expected experimental error up to 15%. Binding isotherms are shown to the right of sensorgrams. Family-wide residue numbering is followed by UniProt numbering (in curly brackets) for each mutant. AUC data presented as the mean of two independent measurements, with errors ± the difference of each of these from the mean.

C

| Protein | Oligomeric state | $K_D$ dimerization (µM) |
|---|---|---|
| DIP-α K10Q {K81Q} | dimer | 19.7 ± 1.9 |
| DIP-α K10Q G4S {K81Q G74S} | dimer | 46.3 ± 5.7 |
| DIP-α K10Q S32D {K81Q S133D} | dimer | 64.4 ± 3.9 |

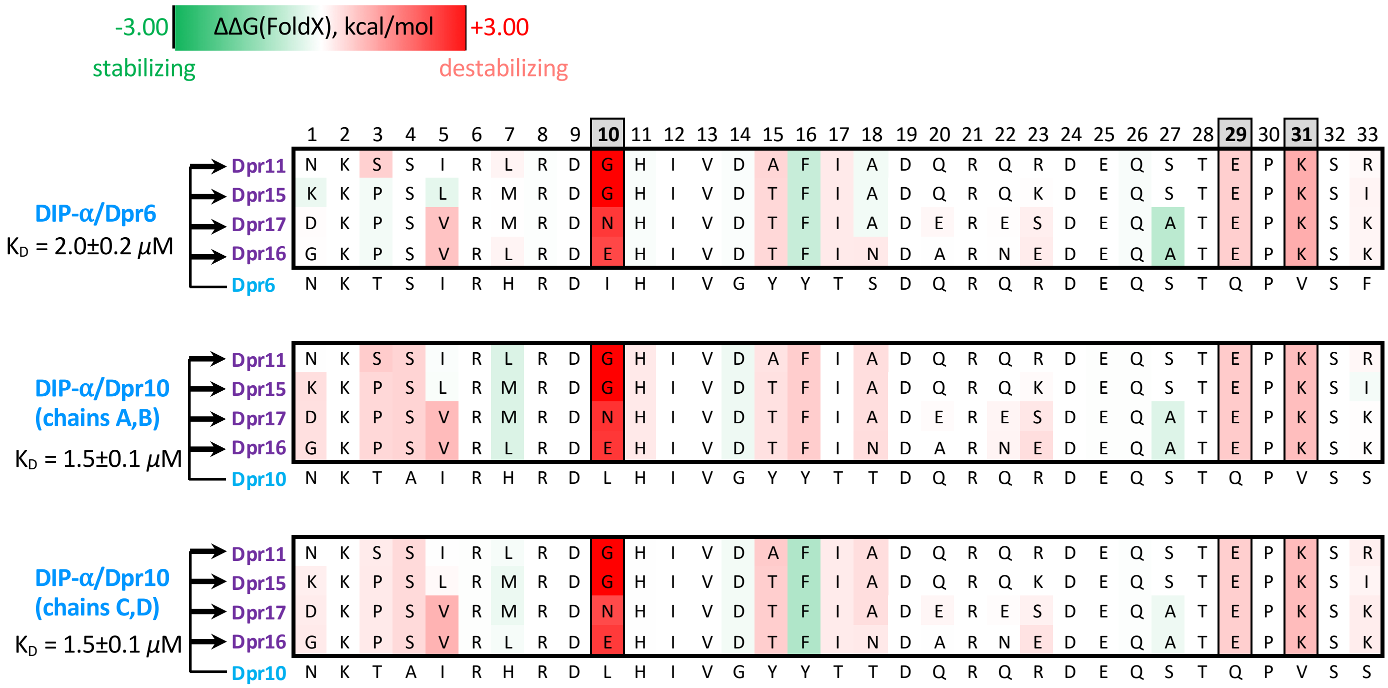


**Supplementary Fig. 2:** Extended version of Fig. 3A. Negative constraints on the purple Dpr surface, which prevent inter-subgroup binding of the four purple group Dprs with DIP-α are assigned based on consensus of predictions using three representative blue group complexes: DIP-α/Dpr6, and the two distinct conformations observed in crystals of DIP-α/Dpr10. Positions that pass both energy and evolutionary filters are shown in grey. All notations as in Fig. 3A. Average $K_D$ values ± standard deviation are given for each interaction based on a number of independent SPR experiments, see methods.

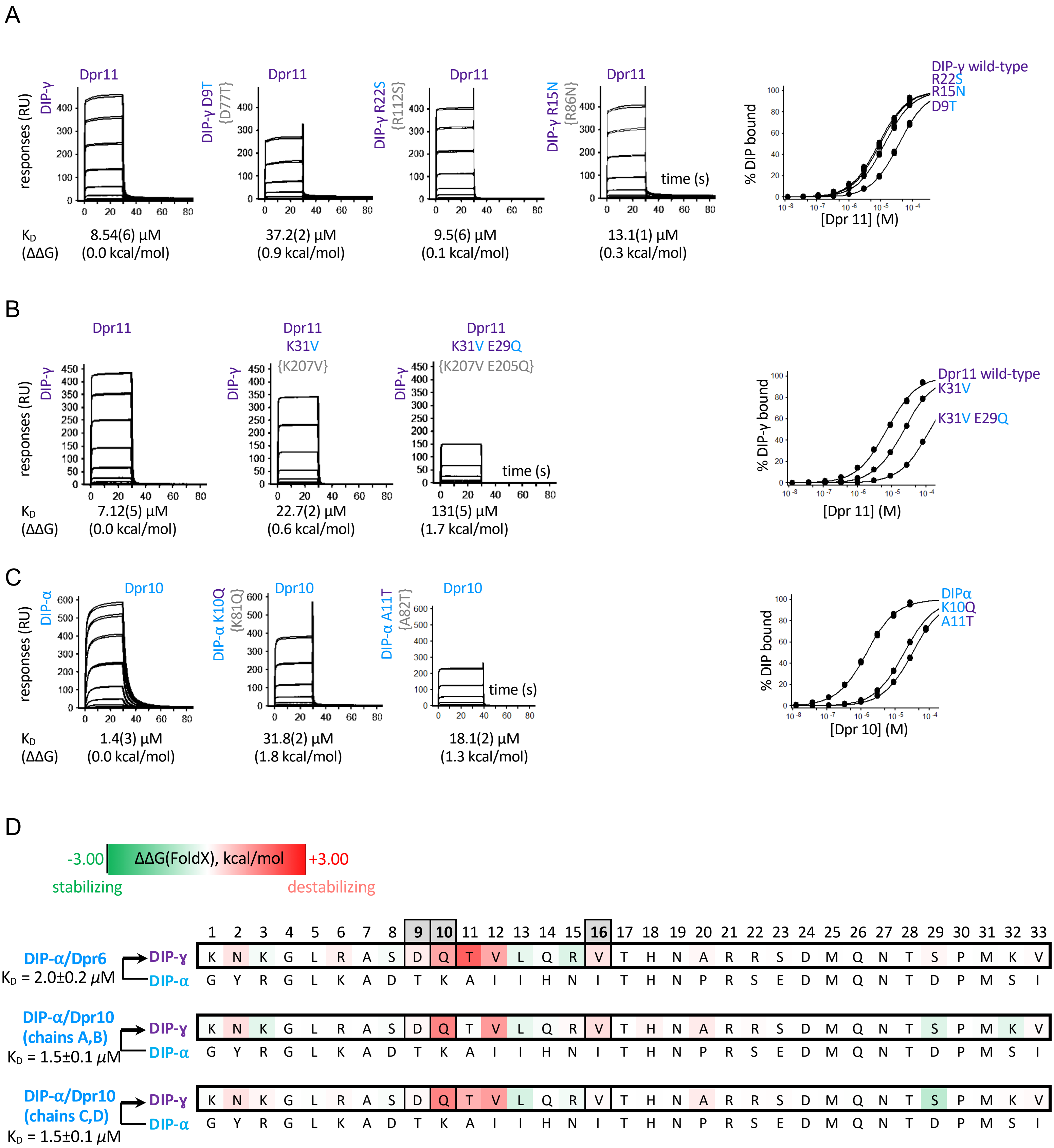


**Supplementary Fig. 3: SPR experiments supporting prediction of negative constraints that prevent blue to purple inter-subgroup binding on (A) DIP-α surface, (B) blue Dpr surface, (C), DIP-γ surface.** All notations as in Fig. 3C. Each row shows SPR sensograms of Dpr analytes binding over individual DIP-immobilized surfaces. An overlay of the binding isotherms for each surface is shown to the right. $K_D$ (ΔΔG) values for each DIP/Dpr interaction are listed below the SPR sensorgrams. For each $K_D$, the number in parenthesis represents the fitting error in the last significant figure, in μM, for a single experiment with an expected experimental error up to 15%. Family-wide numbering of interfacial positions is used for all the mutants. Uniprot numbering is given in curly brackets. **(D).** Extended version of Fig. 4C. Negative constraints on the DIP-γ surface, which prevent inter-subgroup binding to the blue group Dprs are assigned based on consensus of predictions using three representative blue group complexes: DIP-α/Dpr6, and the two distinct conformations observed in crystals of DIP-α/Dpr10. All notations as in Fig. 3A. Average $K_D$ values ± standard deviation are given for each interaction based on a number of independent SPR experiments, see methods.

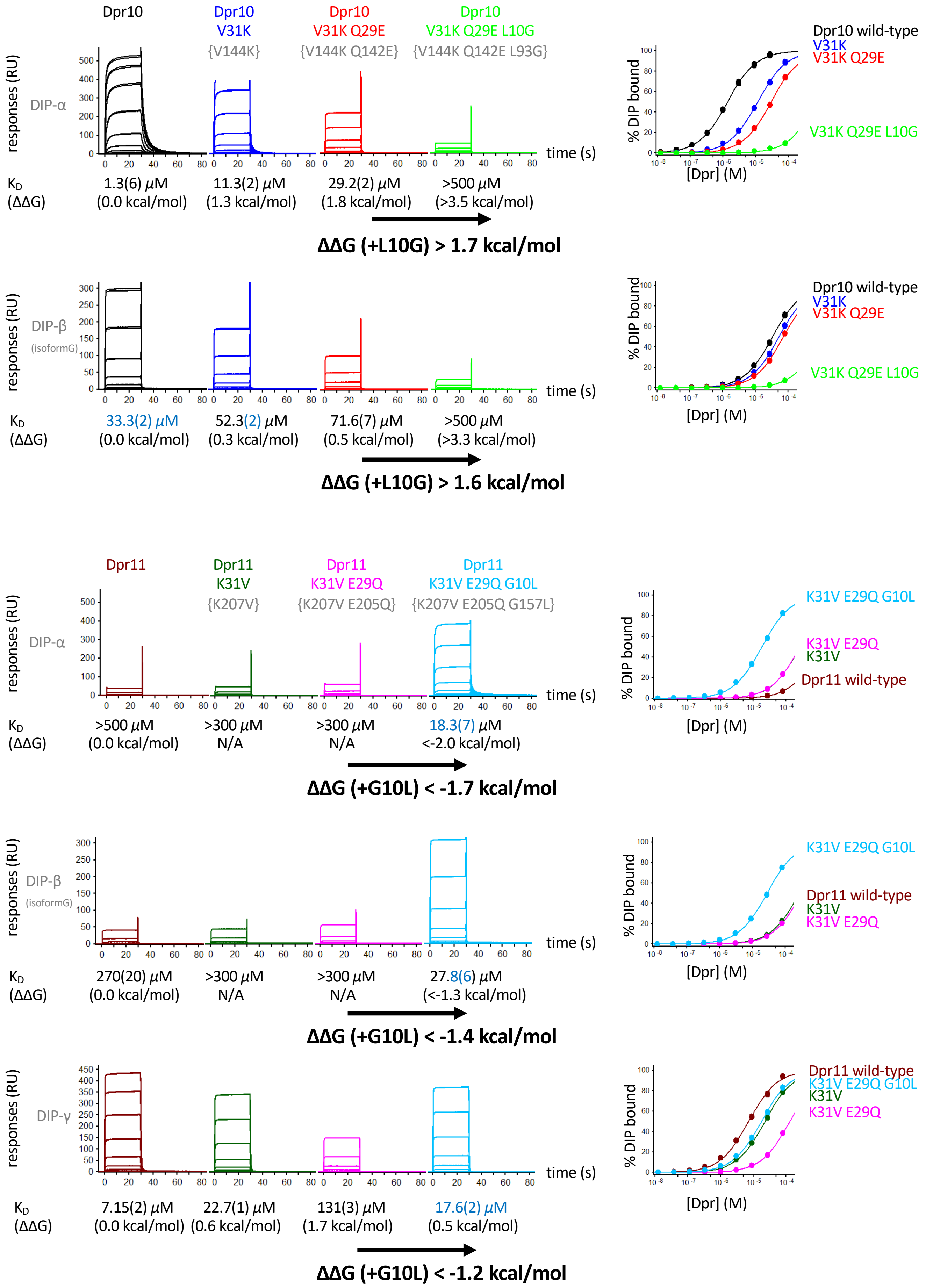


**Supplementary Fig. 4: SPR experiments of the binding of Dpr10, Dpr11, and their mutants to DIP-α, DIP-β, and DIP-γ.** Each row shows SPR sensorgrams of Dpr analytes binding over an individual DIP-immobilized surface. Binding isotherms of DIP/Dpr interactions are given to the right of SPR sensorgrams. $K_D$ values are given under each sensorgram. The number in parenthesis represents the fitting error in the last significant figure, in μM, for a single experiment with an expected experimental error up to 15%. Family-wide numbering of interfacial positions is used for all the mutants. Uniprot numbering is given in curly brackets.
